# EPP1 is an ancestral component of the plant Common Symbiosis Pathway

**DOI:** 10.1101/2025.09.30.679610

**Authors:** Mélanie Rich, Tatiana Vernié, Manish Tiwari, Lucie Chauderon, Julie Causse, Tifenn Pellen, Alice Boussaroque, Matheus E. Bianconi, Michiel Vandenbussche, Pierre Chambrier, Aurélie Le Ru, Baptiste Castel, Sairam Nagalla, Julie Cullimore, Jean Keller, Oswaldo Valdes-Lopez, Malick Mbengue, Jean-Michel Ané, Pierre-Marc Delaux

## Abstract

The success of plants on land has been enabled by mutualistic intracellular associations with microbes for 450 million years (Delaux and Schornack 2021). Because of their intracellular nature, the establishment of these interactions requires tight regulation by the host plants. In particular, three genes – *SYMRK, CCaMK* and *CYCLOPS* – form the core of an ancestral common symbiosis pathway (CSP) for intracellular symbioses, and are conserved since the most recent common ancestor of land plants (Radhakrishnan et al. 2020; Delaux et al. 2015; Wang et al. 2010; Parniske 2008). Here, we describe *EPP1* as a fourth gene committed to the CSP. Among land plants, *EPP1* is conserved only in species able to associate with at least one type of intracellular symbiont. We found that loss-of-function *epp1* mutants or *EPP1* knock-down lines in four clades of land plants – legumes, Solanaceae, monocots and bryophytes – are all impaired in their ability to associate with arbuscular mycorrhizal fungi. We discovered that the plasma membrane-localized receptor-like SYMRK phosphorylates EPP1 on a conserved serine residue and that this phosphorylation is essential for symbiosis. Using a gain-of-function approach, we demonstrate that EPP1 is upstream of the nuclear kinase CCaMK. We propose that EPP1 is an ancestral component of the essential pathway that has regulated plant symbiosis for half a billion years.

## Main

To successfully colonize lands 450 million years ago, the ancestors of extant land plants had to adapt to a multitude of challenges. Among the innovations that allowed this adaptation, the arbuscular mycorrhizal (AM) symbiosis formed with Glomeromycota fungi facilitated the uptake of water and nutrients which were scarce in this new environment (Beerling 2007). Following the evolution of this first intracellular symbiosis, plants have diversified their symbiotic options with the evolution of alternative interactions such as the ericoid mycorrhizal symbiosis, or the root-nodule bacterial nitrogen-fixing (RN) symbiosis occurring in a monophyletic group of four angiosperm orders – including legumes such as soybean or pea (Radhakrishnan et al. 2020). Forward genetics in model legumes identified a unique signaling pathway – the Common Symbiosis Pathway (CSP) – triggered by the perception of symbiont-derived signals through various plasma-membrane localized LysM-RLK receptors, essential for the AM and RN symbioses (Parniske 2008; Zhang et al. 2024). Perception of the symbiotic signals leads to the phosphorylation of the plasma membrane-localized receptor-like kinase SYMRK (Abel et al. 2024). Following that event, rapid and regular calcium oscillations mediated by the nuclear envelope-localized ion channels DMI1 and CNGCs occur within and around the nucleus (Cook et al. 2025; Charpentier et al. 2016; Ané et al. 2004). These calcium spikes are decoded by the Calcium and Calmodulin dependent Kinase CCaMK (Lévy et al. 2004; Tirichine et al. 2006), leading to its activation and the trans-phosphorylation of the transcription factor CYCLOPS (Messinese et al. 2007; Yano et al. 2008; Singh et al. 2014; Horváth et al. 2011). In legumes, mutants affected in any of these CSP components are unable to form both the AM and the RN symbioses (Parniske 2008).

All the known components of the CSP are deeply rooted in the plant lineage. They are found in the genome of both extant vascular plants and bryophytes, such as the liverwort *Marchantia paleacea*, indicating that the CSP was already present in the most recent common ancestor of the land plants (Wang et al. 2010; Delaux et al. 2015; Radhakrishnan et al. 2020). Knock-out mutants in *M. paleacea* of any CSP member, or of the upstream LysM-RLK receptor, loose symbiotic ability in that species, mirroring the phenotypes observed in angiosperms (Vernié et al. 2025; Teyssier et al. 2025; Lam et al. 2024). This demonstrated that the CSP has had a conserved function in symbiotic signaling for 450 million years. Furthermore, the three core members of the CSP – SYMRK, CCaMK and CYCLOPS – are exclusively conserved in plant species able to accommodate intracellular symbionts, and convergently lost in plant lineages unable to form endosymbioses such as the Brassicaceae in angiosperms, the water fern Azolla, or the liverwort *Marchantia polymorpha* (Delaux et al. 2014; Bravo et al. 2016; Radhakrishnan et al. 2020; Li et al. 2018). This phylogenetic pattern indicates that intracellular symbiosis is the only selective driver maintaining the CSP across land plants (Radhakrishnan et al. 2020). Since the discovery of the CSP in legumes in the early 2000s, the mechanisms by which the signal is transduced from plasma-membrane signaling events leading to SYMRK phosphorylation, to the activation of calcium spiking and its decoding in the nucleus by CCaMK has remained unknown. Quantitative phosphoproteomics in response to symbiotic signals in *Medicago truncatula* revealed dozens of phosphorylation events within 30 minutes (Rose et al. 2012), including on Ser-77 of Early Phosphorylated Protein 1 (EPP1, Valdés-López et al. 2019). Chimeric expression of a silencing construct targeting two *EPP1* paralogs (MtrunA17_Chr3g0132211 and MtrunA17_Chr2g0302244) in transgenic roots resulted in diminished calcium spiking and RN symbiosis (Valdés-López et al. 2019). In addition, expression of a phosphomimic version of EPP1, *MtEPP1*^*S77D*^, led to the activation of a number of symbiosis-responsive genes (Ferrer-Orgaz et al. 2024). Altogether, this suggested that EPP1 might contribute to symbiotic signaling, although the presence of three *EPP1* paralogs in *M. truncatula* did not allow further testing its symbiotic function.

Here, we employed a combination of phylogenomics, genetics in four different plant clades, biochemistry and epistasis assays to decipher the role of EPP1. We conclude that EPP1 constitutes the fourth component committed to the CSP, filling a gap between plasma membrane and nuclear symbiotic signaling events.

### Conservation of EPP1 across land plants

A unifying feature of the CSP-dedicated members is their strict conservation across land plants, and their losses in lineages that have lost the ability to host intracellular symbionts (Radhakrishnan et al. 2020). To determine whether *EPP1* follows this pattern, we gathered a set of 247 genomes covering all the major land plants clades and diverse symbiotic abilities (Figure 1, Supplementary Table 1), including basal angiosperms (1 intracellular symbiosis-forming species / 2 non-symbiotic forming ones), dicots (135/26), magnoliids (5/0), monocots (13/11), gymnosperms (7/2), ferns (3/4), lycophytes (5/2), mosses (1/6), hornworts (11/2) and liverworts (4/5). We queried this dataset for *EPP1* homologs and reconstructed its phylogeny (Figure 1, Supplementary Figure 1). For each clade we determined the conservation rate of EPP1 in species able to accommodate intracellular symbionts and in those which have lost this ability (Supplementary Table 1). *EPP1* was detected in most species forming intracellular symbioses, in both the vascular plants (94%) and the bryophytes (95%, Supplementary Table 1, Figure 1). In contrast, *EPP1* was absent in all species that have lost the ability to accommodate intracellular symbionts, irrespective of their phylogenetic position (Figure 1), with the two exceptions being the angiosperm *Cephalotus japonicus* which also retains other members of the CSP (Radhakrishnan et al. 2020), and the hornwort *Megaceros flagellaris*. In particular, and similar to the other three committed members of the CSP, plant species that establish tight, but not intracellular symbioses, do not retain *EPP1* (Supplementary Table 1). This includes for example the water fern Azolla which associates with cyanobacteria, or the Gymnosperm Pinus which forms an ectomycorrhizal symbiosis (Supplementary Table 1).

**Figure 1.**
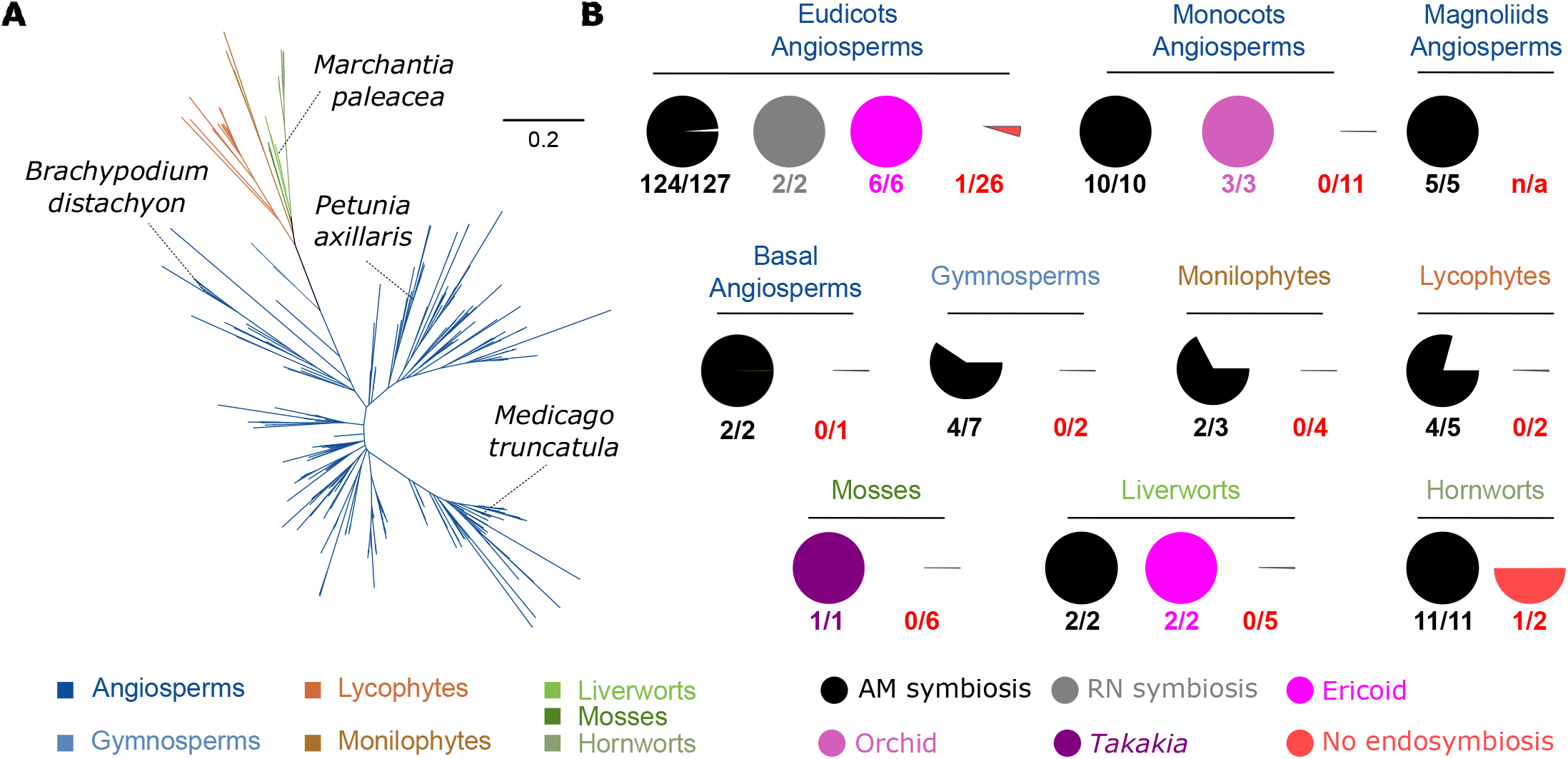
*EPP1* displays a symbiosis-associated phylogenetic pattern. **A**. Maximum-likelihood phylogenetic tree of *EPP1* indicating its occurrence in the bryophyte and vascular plant lineages. **B**. Conservation pattern of *EPP1* depending on the symbiotic abilities of the plant species. For each plant clade, the colored discs indicate the retention rate for species exclusively forming one or another type of endosymbiosis (black, grey, magenta, purple, and pale pink) or not able to form endosymbiosis (red).

The very specific distribution of *EPP1* across land plants therefore indicates that intracellular symbiosis has been the main selection driver maintaining it in plant genomes for 450 million years.

### EPP1 is essential for symbiosis

To define the role of *EPP1* in intracellular symbiosis, we used stable lines in the legume *Medicago truncatula* expressing an RNAi construct targeting two *EPP1* paralogs (Ferrer-Orgaz *et al*. 2024). The resulting MtEPP1^RNAi^ lines showed *EPP1* transcripts down-regulation ranging from 62% to 78%. Quantification of the mycorrhization rate 7 weeks after inoculation with the model AM fungus *Rhizophagus irregularis* showed impaired colonization in these lines and reduced arbuscule formation (Figure 2A). While wild-type *M. truncatula* plants were colonized at 41%, this was reduced to 31–32% in the EPP1^RNAi^ lines. Together with previously published data showing aborted RN symbiosis in *M. truncatula* stable RNAi lines (Ferrer-Orgaz *et al*. 2024) and chimeric transgenic roots expressing an RNAi construct targeting *EPP1* (Valdés-López et al. 2019), this indicates that *EPP1* is required for both types of intracellular symbioses in legumes.

**Figure 2.**
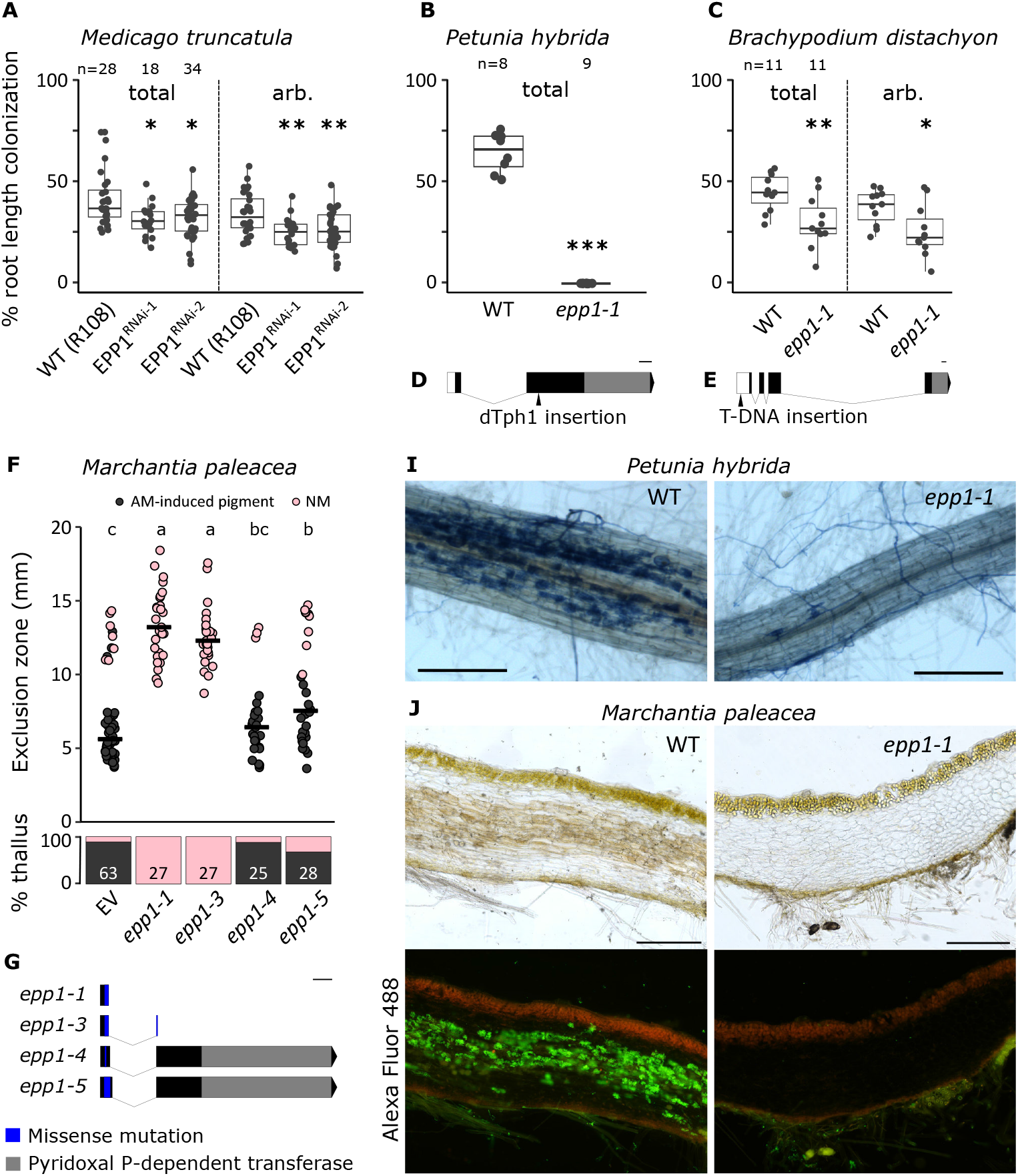
EPP1 is necessary for arbuscular mycorrhiza symbiosis. **A-C** Percentage root length colonization of *Medicago truncatula* EPP1^RNAi^ lines (**A**), *Petunia hybrida* (**B**) and *Brachypodium distachyon* (**C**) mutant lines. Stars indicate significant differences between wild-type and mutants (Welch t-test). **F** Colonization quantification of *Marchantia paleacea epp1* mutants with distance between the thallus notch and colonized area (top) and percentage of colonized thalli (bottom). Bar represent median value. Number of observed thalli is indicated. Letters indicate statistical differences (ANOVA, posthoc Tukey). Each bar corresponds to an independent transformed line. NM = Non-Mycorrhized. **DEG** Schematic representations of the *epp1* mutations in *P*.*hybrida* (**D**), *B*.*distachyon* (**E**) and *M. paleacea* (**G**). Scale bar = 100bp. **IJ** Representative pictures of the phenotypes of *P. hybrida* (**I**) and *M. paleacea* (**J**) *epp1* and their respective wild types. *R. irregularis* is visualized with blue ink (**I**) or with WGA-Alexa Fluor 488 (**J**). *P. hybrida* images were taken using a Zeiss Axiozoom V16 microscope, *M. paleacea* using a Nikon Ti Eclipse. Scale bar = 0.25 mm (**I**), 0.5mm (**J**).

Given its conservation across land plants, we hypothesized that *EPP1* would be required for the widespread AM symbiosis beyond legumes. To test this hypothesis, we isolated a transposon-insertion mutant in the single *EPP1* of the non-legume dicot *Petunia hybrida* (Solanaceae). *P. hybrida epp1* mutant and its wild-type sibling were inoculated with spores of *R. irregularis*, and mycorrhizal colonization was quantified 5 weeks after inoculation. The wild-type siblings displayed an average colonization rate of 63% (Figure 2B), with the presence of intracellular hyphae, as well as arbuscules, the structure allowing symbiotic nutrient exchanges (Figure 2I). In contrast, the *Phepp1-1* mutant displayed a complete absence of colonization, with only extracellular hyphae surrounding the root, and the formation of aborted hyphopodia on the root surface (Figure 2B, 2I).

Among vascular plants, the split between monocots and dicots is estimated to be as old as 162 million years (Li et al. 2019). To further test the conserved function of *EPP1* across land plants, we isolated a T-DNA insertion line in the *EPP1* ortholog of the monocot *Brachypodium distachyon*. The insertion was located in the promoter region of *BdEPP1*, 179 bp upstream of the start codon, resulting in 90% reduction in gene expression (Supplementary Figure 2). While wild-type *B. distachyon* showed 44% of colonization 6 weeks after inoculation with *R. irregularis*, the *Bdepp1-1* line displayed reduced colonization (30%) and arbuscule formation (Figure 2C). Altogether these results demonstrate that EPP1 is essential for intracellular symbiosis across angiosperms.

The vascular plants and bryophytes lineages last shared a common ancestor 450 million years ago. Among bryophytes, the liverwort *Marchantia paleacea* has become a model species for investigating the conservation of symbiotic processes (Rich et al. 2021; Kodama et al. 2022; Vernié et al. 2025; Teyssier et al. 2025; Sgroi et al. 2024). Using CRISPR/Cas9, we generated seven *M. paleacea epp1* mutants (Supplementary Figure 3). *M. paleacea* transformed with an empty vector and the *M. paleacea epp1* mutant alleles were inoculated with *R. irregularis* and harvested five weeks later. After tissue clearing, the pigment known to accumulate in *M. paleacea* upon colonization by *R. irregularis* (Kodama et al. 2022) was clearly visible in most of the empty-vector controls (89%, Figure 2F, J). Observation by fluorescent microscopy of WGA-Alexa Fluor 488-stained sections of these plants revealed the presence of intracellular hyphae and arbuscules (Figure 2J). By contrast, the *epp1* null alleles showed almost no signs of AM symbiosis-induced pigment, with *epp1-1* showing colonization event only in 4 out of 212 total observed apexes (Figure 2F, J, Supplementary Figure 3). Expression of *MpaEPP1* under its native promoter (*MpaEPP1*_*pro*_) fully complemented AM establishment in the *M. paleacea epp1-1* mutant, further demonstrating the causal link between symbiosis defect and the loss of *EPP1* functions in the mutants (Figure 3B and 4B).

**Figure 3.**
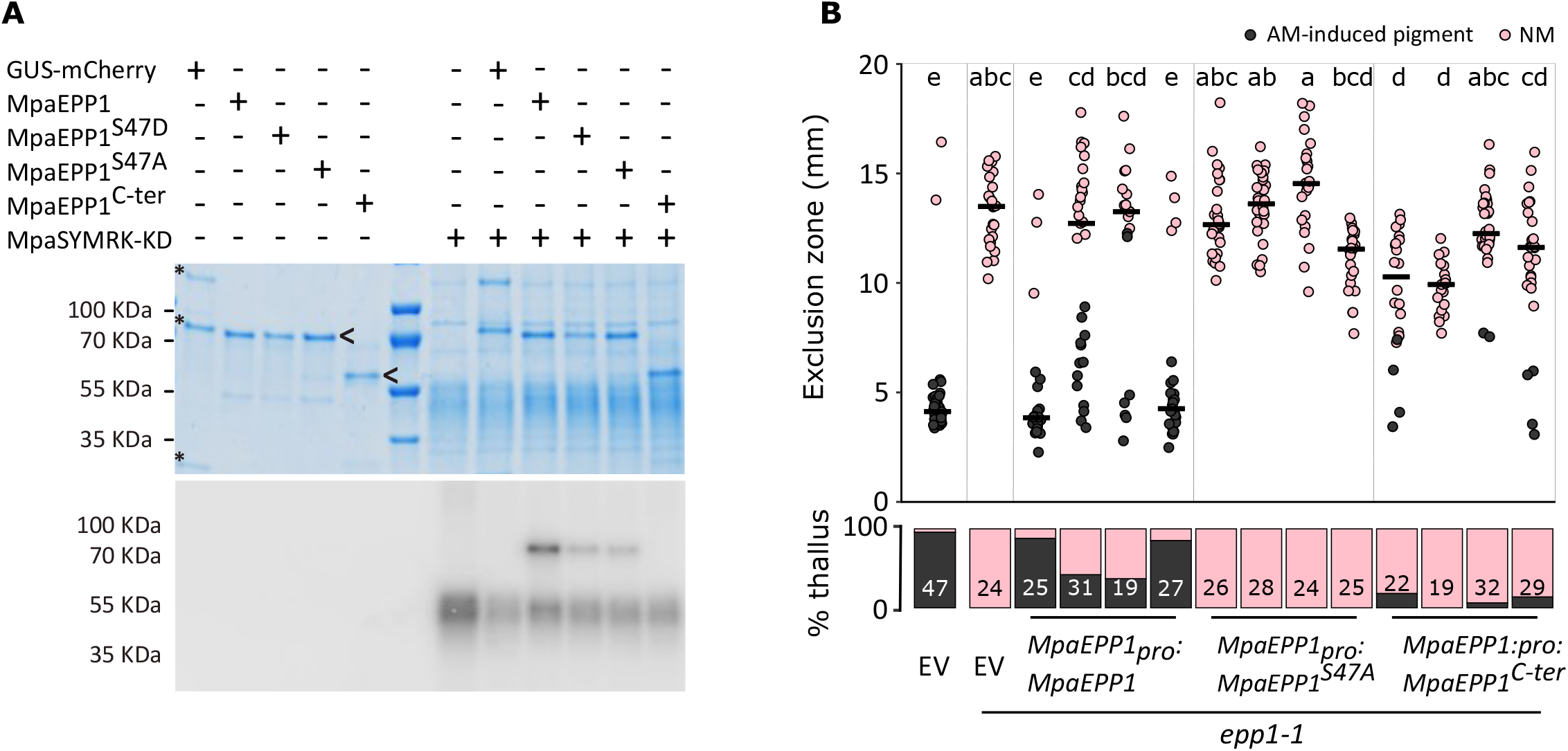
MpaEPP1 serine 47 is a target of SYMRK phosphorylation. **A** Transphosphorylation of MpaEPP1 by the MpaSYMRK kinase domain. *E. coli* expressed SYMRK-KD was used in ^32^P-ATP kinase assays using mCherry fused EPP1 or GUS expressed in *Nicotiana benthamiana* leaves. Coomassie staining (top) and phosphor-image (bottom). **B** Colonization quantification of *Marchantia paleacea* wild-type plants transformed with an empty vector (EV) or *epp1-1* plants complemented with an empty vector (EV) or wild-type MpaEPP1, MpaEPP1^S47A^ or the C terminal Pyridoxal P-dependent transferase domain of MpaEPP1 under the native *MpaEPP1* promoter. Distance between the thallus notch and colonized area (top) and percentage of colonized thalli (bottom) are shown. Bar represent median value. Number of observed thalli is indicated. Letters indicate statistical differences (ANOVA, posthoc Tukey). Each bar corresponds to an independent transformed line. NM = Non-Mycorrhized.

**Figure 4.**
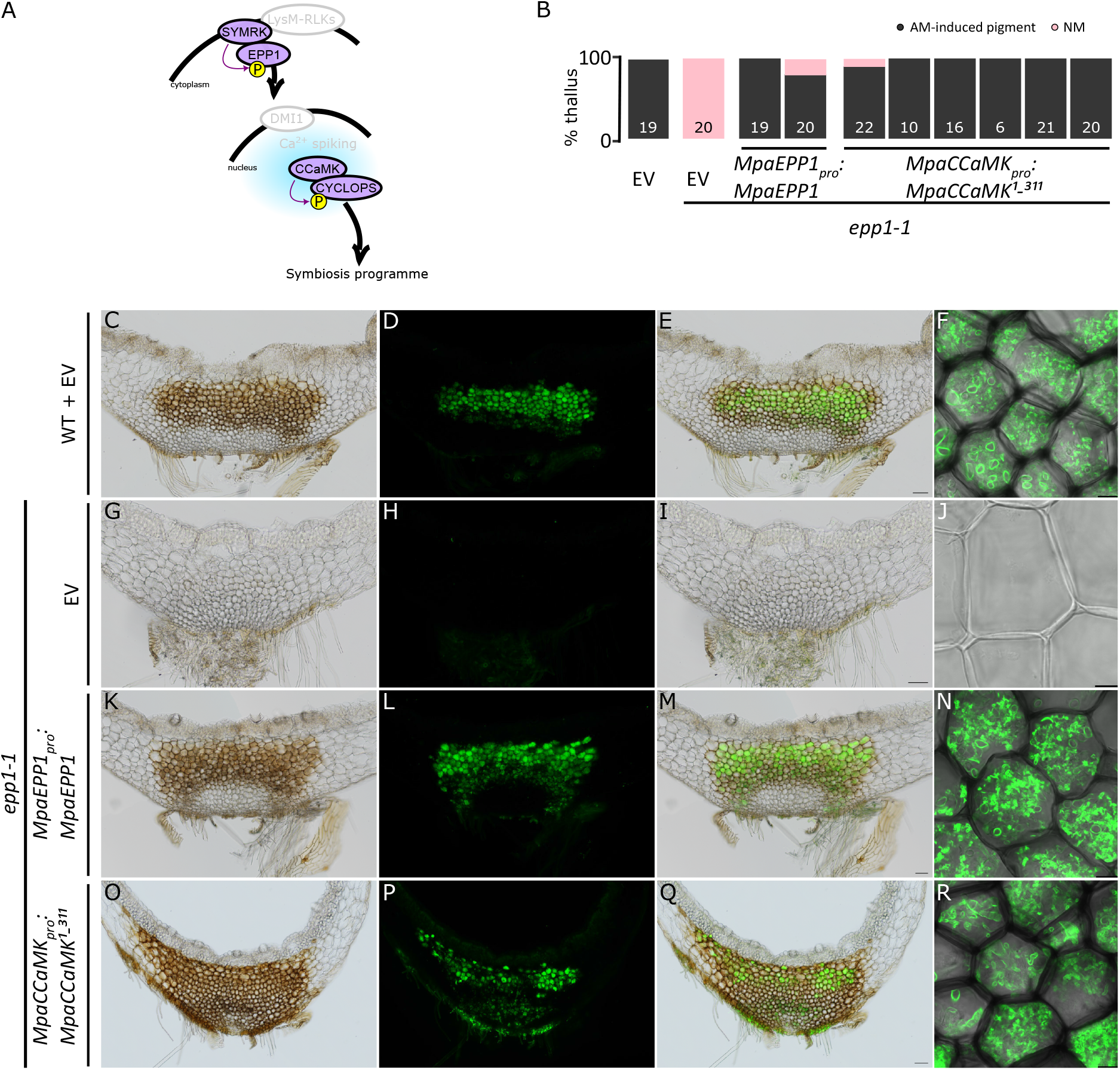
*epp1* AM phenotype is complemented by the autoactive gain of function MpaCCaMK^1-311^ in *Marchantia paleacea*. **A** Proposed model for common symbiotic pathway (CSP) in land plants. **B** Percentage of colonized thalli in wild type plants transformed with an empty vector (EV) or *epp1-1* plants complemented with MpaEPP1 under the native *MpaEPP1* promoter or MpaCCaMK^1-311^ under the native *MpaCCaMK* promoter or an empty vector (EV). Number of observed thalli are indicated. Each bar corresponds to an independent transformed line. NM = Non-Mycorrhized. **C-R** transversal sections of wild-type line transformed with an empty vector (**C-F**) and ***epp1-1*** line transformed with an empty vector (**G-J**) or MpaEPP1 under *MpaEPP1* native promoter (**K-N**) or MpaCCaMK^1-311^ under *MpaCCaMK* native promoter (**O-R**) six weeks post inoculation with *R. irregularis. R. irregularis* is visualized with WGA-Alexa Fluor 488. Close-up views of cells in the colonized area by confocal microscopy (**F, J, N, R**). Bright field (**C, G, K, O**), Alexa Fluor 488 (**D, H, L, P**), and overlays (**E, I, M, Q**) are shown for each line. Scale bar: 100µm (Nikon microscopy and 10µm (confocal microscopy).

Altogether these data demonstrate that EPP1, like the other three committed members of the CSP – SYMRK, CCaMK and CYCLOPS – has had an essential function in arbuscular mycorrhizal symbiosis since the most recent common ancestors of the land plants.

### SYMRK phosphorylates EPP1

Previous analyses in legumes suggested that EPP1 might contribute to symbiotic signaling upstream of calcium spiking (Ferrer-Orgaz et al. 2024; Valdés-López et al. 2019). Given its cytosolic localization and phosphorylation in response to symbiotic signals in legumes (Ferrer-Orgaz et al. 2024; Valdés-López et al. 2019), we hypothesized that EPP1 could be a substrate of the SYMRK kinase. To test this hypothesis, we produced recombinant *MpaEPP1* fused to mCherry in *N. benthamiana* leaves. The tagged protein was produced at the expected size, and no cleavage was observed (Supplementary Figure 4). In addition to the full-length MpaEPP1-mCherry, we produced an mCherry-tagged version of MpaEPP1 lacking the conserved N-terminal domain but retaining the domain annotated as a PLP-dependent transferase (MpaEPP1^C-ter^). These different versions were incubated with purified recombinant GST-MpaSYMRK-KD kinase domain to test for trans-phosphorylation. GST-MpaSYMRK-KD autophosphorylates, but in addition trans-phosphorylation of MpaEPP1-mCherry was clearly detected (Figure 3A), while MpaEPP1^C-ter^-mCherry or a GUS-mCherry control did not produce any signal (Figure 3A). In legumes, phosphorylation in response to symbiotic signals occurs on Ser-77 in *M. truncatula* (Valdés-López et al. 2019). This Serine residue is highly conserved (Supplementary Dataset 1), irrespective of the type of intracellular symbiosis formed by the species (Supplementary Dataset 1). Beyond the position in the primary alignment, AlphaFold modeling revealed that this essential Serine residue sits in a short loop between two conserved alpha helixes, suggesting structural constraints on its position (Supplementary Figure 5). We produced MpaEPP1^S47A^- and MpaEPP1^S47D^-mCherry versions in *N. benthamiana*. Both EPP1 variants were stable (Supplementary Figure 4). We then tested the ability of GST- MpaSYMRK-KD to phosphorylate these EPP1 variants. In contrast with MpaEPP1-mCherry, MpaEPP1^S47A^-mCherry and MpaEPP1^S47D^-mCherry displayed only residual phosphorylation (Figure 3A). These results demonstrate that EPP1 is a phosphorylation target of SYMRK, with the highly conserved and structurally constrained Serine within its N-terminal region being a primary phosphorylation site.

To test the relevance of this phosphorylation site in signaling, we conducted complementation assays using the *M. paleacea epp1* mutant and MpaEPP1 variants. The *M. paleacea epp1-1* allele that showed barely any colonization by arbuscular mycorrhizal fungi (Figure 2F, J, Supplementary Figure 3) was retransformed with MpaEPP1^C-ter^ and MpaEPP1^S47A^ under the active *MpaEPP1*_*pro*_ promoter (Figure 3B, Supplementary Figure 6). The resulting lines were then compared to lines retransformed with the wild-type *MpaEPP1* or an empty-vector. *M. paleacea epp1-1* mutant lines transformed with *MpaEPP1*_*pro*_*:MpaEPP1* showed almost full complementation (Figure 3B). By contrast, *M. paleacea epp1-1* retransformed with the empty-vector, *MpaEPP1pro:MpaEPP1*^*C-term*^ or *MpaEPP1pro:MpaEPP1*^*S47A*^ did not show complementation (Figure 3B), defining the phosphorylation of Ser-47 as essential for the symbiotic function of EPP1.

We conclude that the phosphorylation of EPP1 by SYMRK is an essential signaling step for the establishment of the arbuscular mycorrhizal symbiosis. Conservation of the Serine targeted by SYMRK across land plants in EPP1 suggests that this regulation is an ancestral feature of the CSP.

### EPP1 is a bona fide CSP member

The phylogenomic, biochemical and genetic data presented here strongly suggest that EPP1 is a *bona fide* member of the CSP acting between SYMRK and CCaMK. To further test this hypothesis, we expressed a phosphomimic version of *MpaEPP1* (*MpoEF1a*_*pro*_*:MpaEPP1*^*S47D*^) in a *M. paleacea* background expressing a *GUS* reporter under control of the *CYC-RE*_*pro*_ promoter, known to be induced upon activation of CYCLOPS by CCaMK (Singh et al. 2014; Vernié et al. 2025). In these lines, faint, but consistent, GUS signal could be detected in the area of the thallus prone to infection by *R. irregularis* (Supplementary Figure 7). Given that EPP1 is phosphorylated by SYMRK, we reasoned that it might be positioned between SYMRK and the nuclear symbiotic responses. Nuclear calcium-spiking downstream of SYMRK is decoded by CCaMK. Autoactive, Gain-of-Function, versions of CCaMK have been discovered in legumes (Tirichine et al. 2006; Gleason et al. 2006). Expression of one of them, the truncated *CCaMK*^*1-311*^ lacking the autoinhibitory domain, leads to the activation of the symbiotic program in legumes and in *M. paleacea* (Gleason et al. 2006; Tirichine et al. 2006; Vernié et al. 2025). If the function of EPP1 is indeed to act as the link between SYMRK and the activation of CCaMK, we hypothesized that expressing *MpaCCaMK*^*1- 311*^ should complement the symbiotic defect of the *Mpaepp1* mutant. We retransformed the *M. paleacea epp1-1* mutant with an empty vector, or with *MpaCCaMK*^*1-311*^ under control of the functional (Supplementary Figure 8) *MpaCCaMK*_*pro*_ promoter, and inoculated these lines with *R. irregularis*. While none of the empty-vector-retransformed *M. paleacea epp1-1* mutants showed colonization, the *MpaCCaMK*_*pro*_*:MpaCCaMK*^*1-311*^-retransformed ones were fully colonized, displaying intracellular hyphae and arbuscules (Figure 4). The full complementation of *Mpaepp1-1* with *MpaCCaMK*_*pro*_*:MpaCCaMK*^*1-311*^ demonstrates that EPP1 acts upstream of CCaMK in the CSP.

Altogether, these epistasis tests position EPP1 between two core members of the CSP, the plasma membrane-localized receptor-like kinase SYMRK and the nuclear-localized kinase CCaMK.

### Conclusion

Genetics in land plants belonging to four lineages and spanning 450 million years of plant evolution reveals the ancestral role of EPP1 for arbuscule mycorrhizal symbiosis. Previous studies have suggested a similar function for the RN symbiosis in legumes (Ferrer-Orgaz et al. 2024; Valdés-López et al. 2019), and our phylogenetic screen indicates a link with all other types of intracellular symbioses. Phosphorylation of EPP1 by SYMRK and the complementation of symbiosis in the *M. paleacea epp1* mutant by CCaMK^1-311^ positions EPP1 as one of the missing components linking plasma membrane-localized activation of the CSP and the nuclear-localized activation of downstream symbiotic signaling. Altogether, the combination of phylogenetic, reverse genetic, biochemical and epistasis approaches presented here identify EPP1 as a fourth committed member of the ancestral plant Common Symbiosis Pathway.

## Methods

### Phylogenetic analysis

We first used BLAST v2.15.0 (Camacho et al. 2009) to search for putative homologs in 247 genomes encompassing all main lineages of bryophytes and vascular plants, with the protein sequence of *Medicago truncatula EPP1* used as query (MtrunA17Chr3g0132211, e-value = 10-6). All hits were aligned with MAFFT v7.505 (Katoh and Standley 2013) using the automatic mode (--auto, --maxiterate 20), and columns with more than 70% of gaps were removed using trimAl v1.4.1-gt 0.3). A maximum-likelihood tree was then estimated with IQ-TREE v2.2.2.6 (Nguyen et al. 2015) using the standard model selection (-m TEST). This initial tree was inspected to identify the clade of orthologous sequences containing the query. These sequences were then extracted, realigned and trimmed to remove columns with more than 90% of gaps (-gt 0.1). The final phylogenetic tree was estimated as above, with branch support assessed using 1000 pseudoreplicates for the SH-like approximate likelihood test and ultrafast bootstrap.

### Protein structure prediction

The structures of EPP1 from *M. truncatula* (MtEPP1, MtrunA17Chr3g0132211), *Petunia axillaris* (PaEPP1, Peaxi162Scf00518g00423.1), *Brachypodium distachyon* (BdEPP1, Bradi4g19020.1.v3.1) and *Adiantum nelumboides* (AnEPP1, JAKNSL_Ane01050.m), *Marchantia paleacea* (MpaEPP1, utg000002g0001491) were modelled using the AF3 server (Deepmind, 2024).

### Plant material

*M. truncatula* EPP1 (MtrunA17Chr3g0132211) RNAi stable lines L31 (EPP1^RNAi-1^) and L28a (EPP1^RNAi-2^) were described in Ferrer-Orgaz 2023. *P. hybrida epp1-1* (Peaxi162Scf00518g00423) was identified in the sequence-indexed dTph1 transposon database (Vandenbussche et al. 2008). The wild-type sibling was used as a control. *B. distachyon* seeds of the JJ21512 line with a T-DNA insertion 179 bp in front of the ATG of *EPP1* (Bradi4g19020) and the control Bd21.3 were obtained from the Joint Genome Institute and bulked to the T3 generation. *M. paleacea epp1-1* to-*7* (Marpal_utg000002g0001491) were generated using CRISPR/Cas9. *M. paleacea ccamk #9*.*19* and *CYC-RE*_*pro*_*:GUS #5 &10* were described in Vernié et al. 2025.

### Mycorrhization assays

*M. paleacea* mycorrhization assays were performed as described in Vernié et al. 2025. Plants were harvested 5 weeks post inoculation for experiments from Figure 2, 3 and Supplementary Figures 3 and 4 and 6 weeks for experiments in Figure 4. Quantification of colonization was done as described in Kodama et al. 2022.

*P. hybrida* mycorrhization assays were performed as described in Vernié et al. 2025.

For *M. truncatula* mycorrhization assays, seeds were scarified and surface-sterilized as in Valdés-López et al. 2019. Seeds were placed on 1% agar in a cold room for two days, followed by two days in the dark at room temperature and exposed to light until they had the first trifoliate leaf. Seedlings were transferred to round pots (6 cm diameter, 25 cm height) filled with two-thirds level of washed and autoclaved Turface® (E P MINERALS Safe T Sorb 7941) and a one cm layer of fine sand. Plants were inoculated with 200 arbuscular mycorrhizal spores (Mycorise ASP, Premier Tech), and pots were filled up with Turface®. Plants were grown in a growth chamber (conditions: 16h/8h, 23/18°C), fertilized with five ml of Half-Strength Hoagland medium without ammonium phosphate, supplemented with 20μM phosphate (monopotassium phosphate) once a week for the first two weeks and were watered with distilled water daily. Plants were harvested seven weeks after inoculation, roots cut into pieces (1-2cm), rinsed in distilled water, cleared 10% (w/v) KOH at 95°C for 10 minutes, rinced three times in distilled water and stained in Sheaffer ink staining solution at 95°C for 10 minutes. Root colonization was quantified using the grid intersect method.

For *B. distachyon* mycorrhization assays, seeds were sterilized 15 minutes in 2% (v/v) bleach and rinsed (both repeated twice) and soaked for 2 hours. Seeds were placed onto 1% agar for two days in a cold room, followed by two days in the dark at room temperature and a day of light exposure. The seedlings were transferred to pots and were inoculated with 400 arbuscular mycorrhizal spores (Mycorise ASP, Premier Tech). Growth conditions and post-harvesting treatments were similar to those of Medicago.

All experiments were repeated 3 times with similar results

### Cloning and plant transformation

The Golden Gate modular cloning system (Patron et al. 2015) was used to prepare the plasmids as described by Rich et al. 2021. Levels 0, 1, and 2 used in this study are listed in supplementary Table 2 and held for distribution in the ENSA project core collection. Sequences were domesticated (listed in supplementary Table 2), synthesized, and cloned into pMS (GeneArt, Thermo Fisher Scientific, Waltham).

*M. paleacea* transformation was performed as described in Vernié *et al*. 2025.

### GUS assays

GUS assays in *M. paleacea* were performed as described in Vernié *et al*. 2025.

### Microscopy

*M. paleacea* sections were generated similarly to Vernié *et al*. 2025. Images were taken using a Zeiss Axiozoom V16 microscope or a Nikon Ti Eclipse inverted microscope equipped with DS Ri2 camera. Close-up of arbuscules were acquired using a Leica SP8 TCSPC confocal microscope and LAS X software with a 25× water immersion objective (Fluotar VISIR 25×/0.95 WATER).

### Protein expression and kinase assay

The predicted intracellular region of *M. paleacea* SYMRK (Marpal_utg000051g0090241, termed the KD) was cloned into the vector pGEX and the Glutathione-*S*-transferase (GST) tagged protein was expressed in *Escherichia coli* DH5α and the GST-SYMRK-KD fusion protein was purified using glutathione resin (GE Healthcare, USA) as described (Fliegmann et al. 2016). SYMRK-KD was released from the resin using PreScission Protease (GE27-0843-01, Sigma Aldrich, Germany).

EPP1-mCherry (wild-type and variants) and GUS-mCherry fusion constructs were generated via GoldenGate cloning in combination with the *Arabidopsis thaliana UBQ10* promoter and the *Agrobacterium tumefaciens NosT* terminator (Engler et al. 2014). Fusion proteins were separately expressed for 2 days in *Nicotiana benthamiana* leaves *via* agrotransformation prior purification from total soluble proteins extracts using RFP-trap beads (Nano-Trap, Proteintech Group). Briefly, 1 gram of leaf tissues was ground in liquid nitrogen before resuspension in 4 mL cold extraction buffer [Tris-HCl 50 mM pH=7.5, NaCl 100 mM, EDTA 1mM, glycerol 10%] supplemented with 5 mM dithiothreitol (DTT), 0.5% (w/v) polyvinylpolypyrrolidone (PVPP), 1% (v/v) Nonidet (NP-40), 1 mM phenylmethylsulfonyl fluoride (PMSF) and 1% (v/v) plant proteases inhibitor cocktail (Sigma Aldrich P9599). Cell debris was pelleted at 5000g for 10 minutes and supernatants filtered using 20 µm Falcon cells strainers before adding 20 µL of RFP beads per extract. After 3 hours under agitation at 4°C, immunopurified proteins were washed 4 times using 500 µL cold extraction buffer and 4 times using 500 µL storage buffer [Tris-HCl 50 mM pH=7.5, NaCl 100 mM, glycerol 20%]. Resuspended proteins in 30 µL storage buffer were checked for purity via SDS-PAGE and Coomassie staining. Quantities were estimated in comparison to Bovine Serum Albumin (BSA) references.

For kinase assays the proteins were incubated in 10 mM HEPES-HCl pH 7.4, containing 5 mM MgCl_2_, 5 mM MnCl_2_, 20 µM ATP containing 5 µCi ^32^P-ATP at 30°C for 50 min. Reactions were analyzed by SDS-PAGE, followed by Coomassie staining and Phosphor imaging.

### Expression profiling

For RT-qPCR assays, five-day-old control and *epp1-1 B. distachyon* seedlings (n=6) were harvested and RNA was isolated using Zymo’s Direct-zol RNA Miniprep Kit following the manufacturers protocol. The RNA was quantified, and 1μg RNA was used to synthesize cDNA using BioRad iScript™ cDNA Synthesis Kit following the manufacturers protocol. The cDNA was diluted five times, and two μl of the diluted cDNA were used in a 10μl RT-qPCR reaction setup using BioRad SsoAdvanced Universal SYBR Green Supermix. *Actin* (BRADI_1g10630, BdActin_F AGGCCAATCGTGAGAAGATG, BdActin_R AGTCGAGACGGAGGATAGCA) and *GADPH* (BRADI_3g14040v3, BdGAPDH_F ATGGGCAAGATTAAGATCGGAATCAACGG, BdGAPDH_R AGTGGTGCAGCTAGCATTTGAGACAAT) genes were used as endogenous controls for the expression of *EPP1* (BdEPP1_F ATGCGTTTGGTTTTGAGCCA, BdEPP1_R ATTTCAGCATCCCAGAGCCT)

### Data and code availability statements

All data generated in this study are available as part of this manuscript.

## Acknowledgments

We are grateful to the genotoul bioinformatics platform Toulouse Occitanie (Bioinfo Genotoul, https://doi.org/10.15454/1.5572369328961167E12) for providing computing resources. This study was supported by the “Laboratoires d’Excellence (LABEX)” TULIP (ANR-10-LABX-41)” and by the “École Universitaire de Recherche (EUR)” TULIP-GS (ANR-18-EURE-0019). This project was supported by the project Engineering Nitrogen Symbiosis for Africa (ENSA) currently funded through a grant to the University of Cambridge by the Bill & Melinda Gates Foundation (OPP1172165) and the UK Foreign, Commonwealth and Development Office as Engineering Nitrogen Symbiosis for Africa (OPP1172165). This project has received funding from the European Research Council (ERC) under the European Union’s Horizon 2020 research and innovation program (grant agreement No 101001675 - ORIGINS) to P-M.D. MEB has received funding from the European Union’s Horizon 2020 research and innovation program under the Marie Skłodowska-Curie grant agreement No. 101105838 (‘SYMBIOLOSS’). This project was also funded by the United States Department of Energy grant #DE-SC0018247 and a United States Department of Agriculture Hatch grant #WIS05052 to J.M.A. Research at O.V.L laboratory is funded by the Secretaría de Ciencia, Humanidades, Tecnología e Inovación (SECHITI) grant # CBF-2025-I-12.

## Author contributions

Conducted experiment M.K.R., T.V., J.Ca., L.C., T.P., A.L.R., A.B., M.V., P.C., M.M., M.T., O.V.-L., S.N., J.Cu. Contributed resources M.V., B.C. Analyzed the data P.-M.D, M.K.R., T.V., J.Ca., L.C., T.P., M.M., M.T., S.N., J.M.A., J.K, M.E.B, J.Cu. Coordinated the project P.-M.D, J.M.A. Designed the experiments P.-M.D., M.K.R., T.V., J.M.A. M.M., M.T.

P.-M.D. wrote the first draft with contributions from M.K.R, T.V., M.T., S.N., J.M.A., M.M. and O.V.-L. All authors contributed to the final version of the manuscript.

## Extended data figure legends

**Supplementary Figure 1**.

Maximum likelihood phylogenetic tree of EPP1 in land plants. Symbiotic status is shown on the right panel (AMS = arbuscular mycorrhizal symbiosis, RNS = root nodule symbiosis, EcM = ectomycorrhiza). Branch support values are shown next to nodes (SH-like approximate likelihood test/ultrafast bootstrap).

**Supplementary Figure 2**. Relative expression of *EPP1* in *Bbepp1-1*.

Expression of *EPP1* relative to *GAPDH* in 5 days old seedlings of *B. distachyon epp1-1* and WT.

**Supplementary Figure 3**. *M. paleacea epp1* mutants phenotype.

**A** Schematic representation of the *epp1* CRISPR mutations studied in *M. paleacea*. **B** Colonization quantification of *M. paleacea epp1* mutants with distance between the thallus notch and colonized area (top) and percentage of colonized thalli (bottom). Bar represent median value. Number of observed thalli is indicated. Letters indicate statistical differences (ANOVA, posthoc Tukey). **C** Longitudinal sections of inoculated thalli of empty vector control, null allele *epp1-3* and in frame mutant *epp1-7* colored with WGA-Alexa Fluor 488. **D** Transversal sections of empty vector control and a rare infected area of *epp1-1*. Number of observations / total number of thalli out of all experiments is indicated for each genotype. Confocal picture of the arbuscules (right) do not show arbuscule defect.

**Supplementary figure 4**. Full gel pictures of the kinase assay shown in Figure 3.

**Supplementary Figure 5**. Predicted structure of the C-terminal alpha-helixes cluster of EPP1

Alphafold 3 (Deepming, 2024), was used to predict the structure of full length EPP1 from

*Medicago truncatula* (MtEPP1, MtrunA17Chr3g0132211, in yellow, pTM=0.58), *Petunia axillaris* (PaEPP1, Peaxi162Scf00518g00423.1, in green, pTM=0.72), *Brachypodium distachyon* (BdEPP1, Bradi4g19020.1.v3.1, in blue, pTM=0.69) and *Adiantum nelumboides* (AnEPP1, JAKNSL_Ane01050.m, in pink, pTM=0.73), *Marchantia paleacea* (MpaEPP1, utg000002g0001491 in purple, PTM=0.66). This figure only displays the conserved C-terminal alpha-helixes cluster plus 10 amino acids before the first alpha-helix and 10 amino acids after the last alpha-helix. The corresponding amino acids are indicated in the figure. Arrows indicate the conserved serine (shown in red) in the loop between the last and penultimate alpha-helixes of the cluster.

**Supplementary Figure 6**.

Two independent *M. paleacea* transformed lines expressing *MpaEPP1*_*pro*_*:GUS* are shown after staining for GUS activity (Scale bar 1mm) after 6 weeks of inoculation with *R. irregularis* (AM, **B**,**C**,**D**,**F**,**G**,**H**) or without inoculation (control, **A, E**). Bright field (**A, B, E, F**), Alexa Fluor 488 (**C, G**), and overlays (**D**,**H**) are shown for each line. Scale bar: 100µm (Nikon microscopy). Total number of thalli observed are indicated.

**Supplementary Figure 7**.

*M. paleacea* transformed lines expressing *CYC-RE*_*pro:*_*GUS* (two independent lines: *CYC-RE*_*pro*_*:GUS* #5 and 10) or *CYC-RE*_*pro*_*:GUS* + *MpoEF1apro:MpaEPP1* (two independent lines: #2, 3) or *CYC-RE*_*pro*_*:GUS* + *MpoEF1apro:MpaEPP1*^S47D^ (two independent lines: #2, 5) are shown after staining for GUS activity, shown in blue (arrowheads). (Scale bar, 1 mm.) Total number of thalli observed are indicated.

**Supplementary Figure 8**.

**A** Percentage of colonized thalli in wild type plants transformed with an empty vector (EV) or *ccamk #9*.*19* (Vernié *et al*. 2025) plants complemented with MpaCCaMK under the native *MpaCCaMK* promoter or an empry vector (EV). Number of observed thalli are indicated. Each bar corresponds to an independent transformed line. **B-D** transversal sections of *ccamk #9*.*19* line transformed with MpaCCAMK under *MpaCCaMK* native promoter six weeks post inoculation with *R. irregularis. R. irregularis* is visualized with WGA-Alexa Fluor 488. Bright field (**B**), Alexa Fluor 488 (**C**), and overlay (**D**) are shown for one representative line. Scale bar: 100µm (Nikon microscopy).

## Extended data table legends

**Supplementary Table 1**.

Genomic data used for the phylogenetic analysis of EPP1.

**Supplementary Table 2**.

List of GoldenGate constructs used in this study.

**Extended dataset**

**Supplementary Dataset 1**.

Alignment of amino acid sequences of EPP1.

