## Supplementary Figures 1 to 8 for "EPP1 is an ancestral component of the plant Common Symbiosis Pathway"

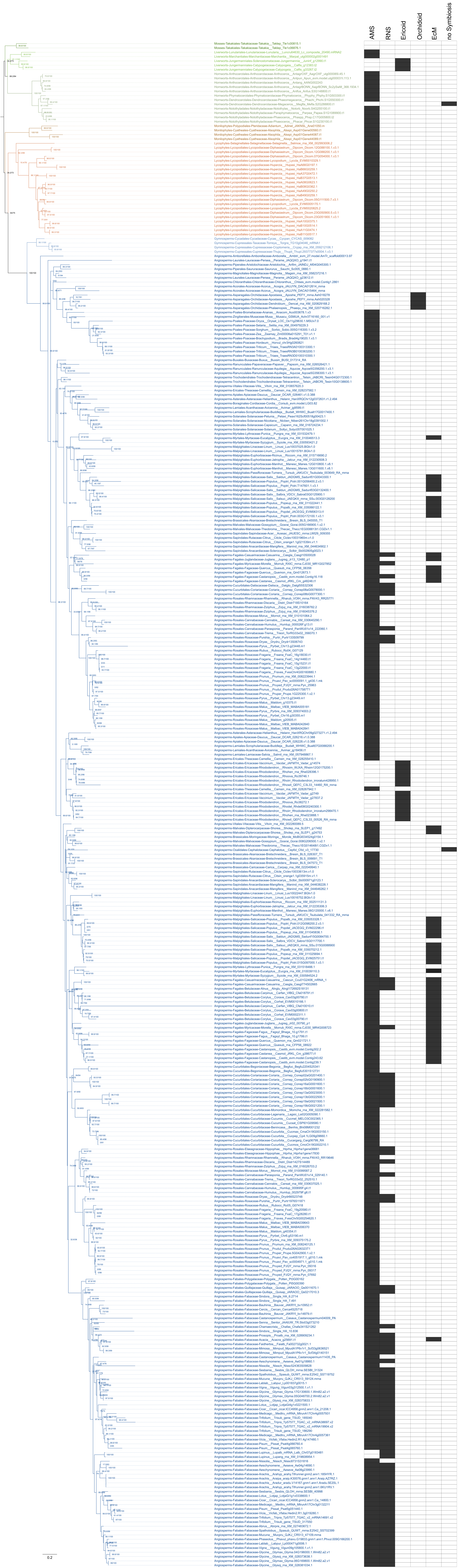
AMS
RNS
Ericoid
Orchidoid
EcM
no Symbiosis
0.2

**Supplementary Figure 1.** Maximum likelihood phylogenetic tree of EPP1 in land plants. Symbiotic status is shown on the right panel (AMS = arbuscular mycorrhizal symbiosis, RNS = root nodule symbiosis, EcM = ectomycorrhiza). Branch support values are shown next to nodes (SH-like approximate likelihood test/ultrafast bootstrap).

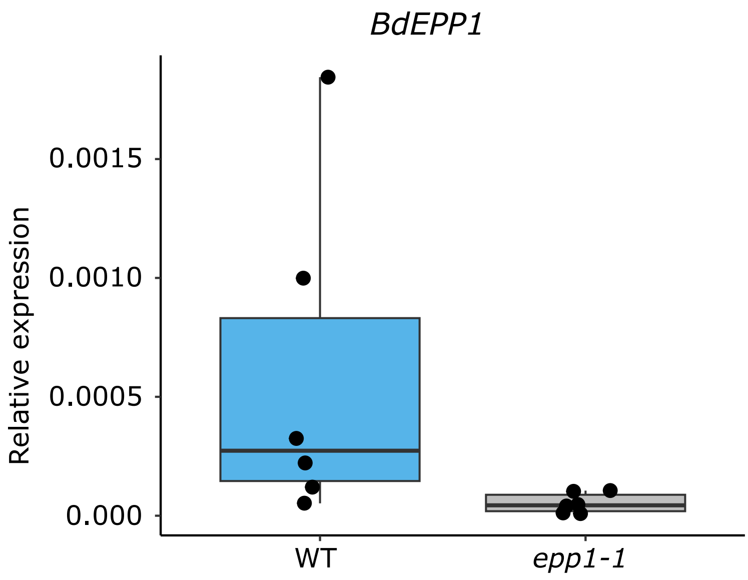


**Supplementary Figure 2.** Relative expression of *EPP1* in *Bbepp1-1.*

Expression of *EPP1* relative to *GAPDH* in 5 days old seedlings of *Brachypodium distachyon epp1-1* and WT

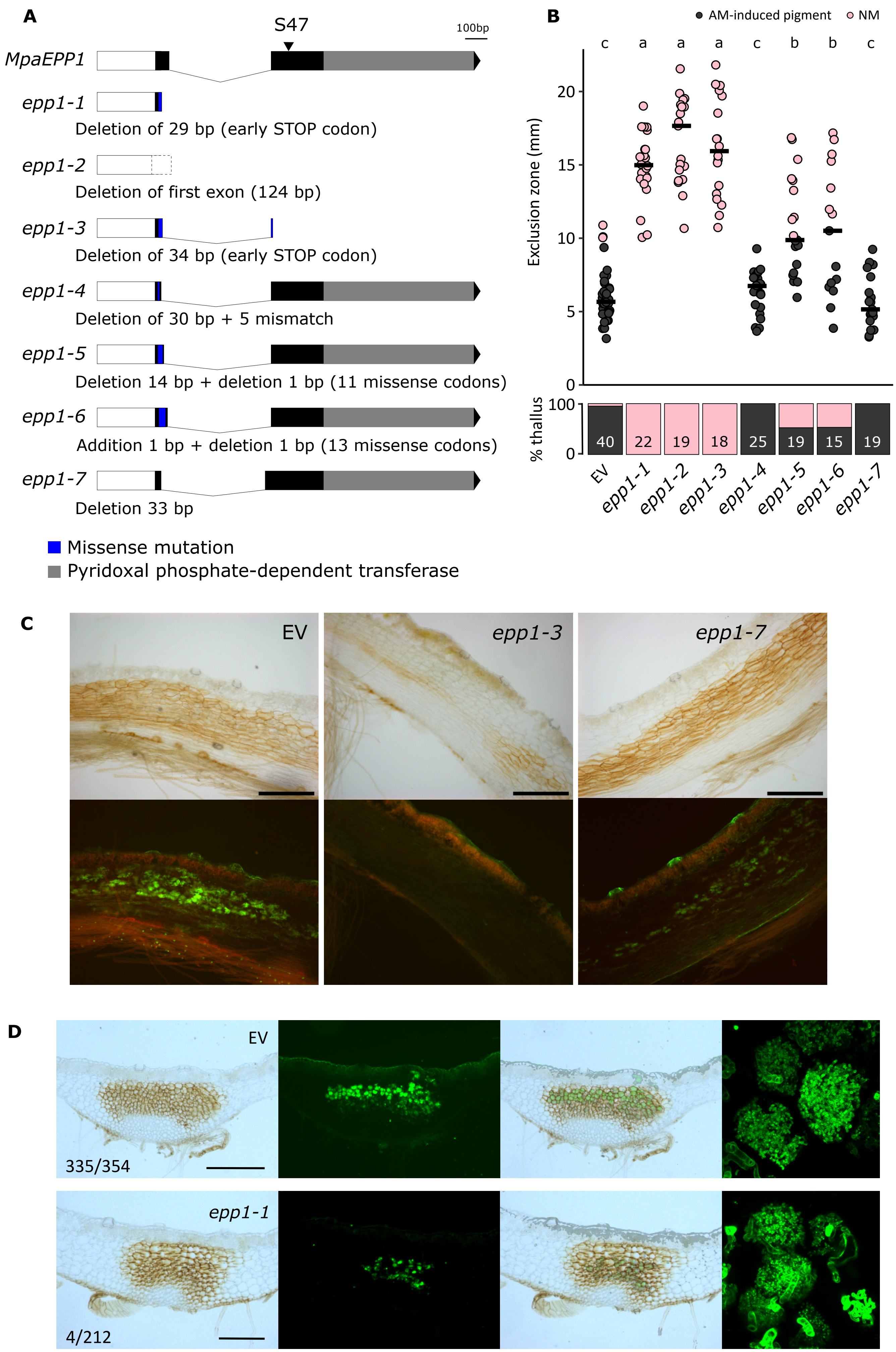

A
S47
100bp
MpaEPP1
epp1-1
Deletion of 29 bp (early STOP codon)
epp1-2
Deletion of first exon (124 bp)
epp1-3
Deletion of 34 bp (early STOP codon)
epp1-4
Deletion of 30 bp + 5 mismatch
epp1-5
Deletion 14 bp + deletion 1 bp (11 missense codons)
epp1-6
Addition 1 bp + deletion 1 bp (13 missense codons)
epp1-7
Deletion 33 bp
Missense mutation
Pyridoxal phosphate-dependent transferase
B
AM-induced pigment
NM
c a a a c b b c
Exclusion zone (mm)
% thallus
40 22 19 18 25 19 15 19
EV epp1-1 epp1-2 epp1-3 epp1-4 epp1-5 epp1-6 epp1-7
C
EV
epp1-3
epp1-7
D
EV
335/354
epp1-1
4/212

**Supplementary Figure 3.** *M. paleacea epp1* mutants phenotype.

**A** Schematic representation of the *epp1* CRISPR mutations studied in *M. paleacea*. **B** Colonization quantification of *M. paleacea epp1* mutants with distance between the thallus notch and colonized area (top) and percentage of colonized thalli (bottom). Bar represent median value. Number of observed thalli is indicated. Letters indicate statistical differences (ANOVA, posthoc Tukey). **C** Longitudinal sections of inoculated thalli of empty vector control, null allele *epp1-3* and in frame mutant *epp1-7* colored with WGA-Alexa Fluor 488. **D** Transversal sections of empty vector control and a rare infected area of *epp1-1*. Number of observations / total number of thalli out of all experiments is indicated for each genotype. Confocal picture of the arbuscules (right) do not show arbuscule defect.

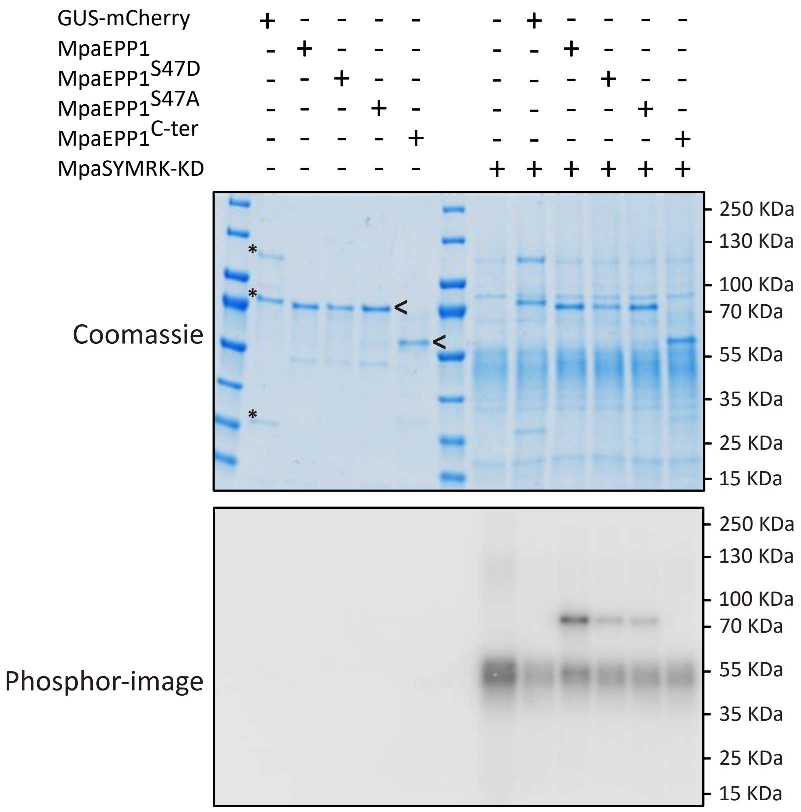


**Supplementary figure 4.** Full gel pictures of the kinase assay shown in Figure 3.

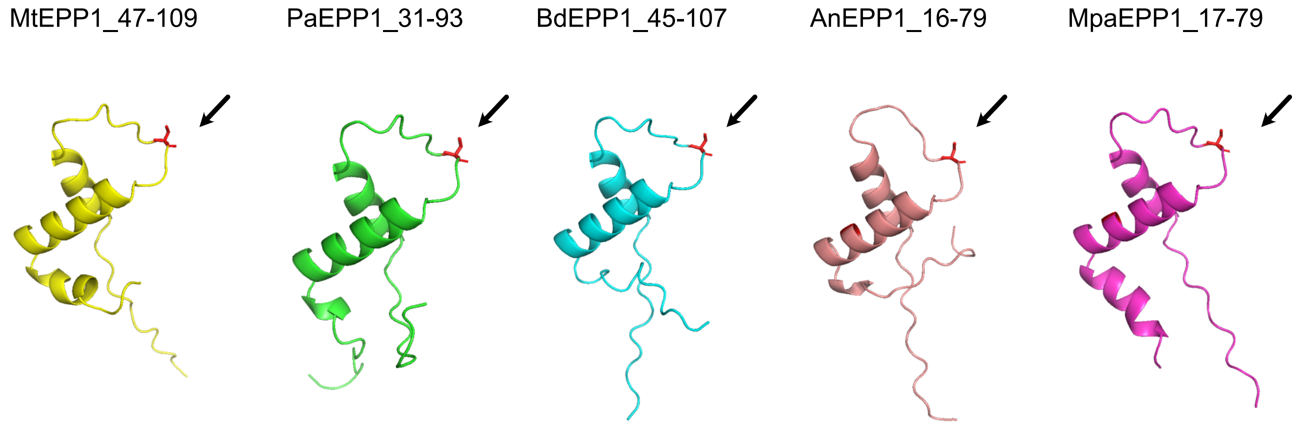


**Supplementary Figure 5**. Predicted structure of the C-terminal alpha-helixes cluster of EPP1.
Alphafold 3 (Deepming, 2024), was used to predict the structure of full length EPP1 from M. truncatula (MtEPP1, MtrunA17Chr3g0132211, in yellow, pTM=0.58), Petunia axillaris (PaEPP1, Peaxi162Scf00518g00423.1, in green, pTM=0.72), Brachypodium distachyon (BdEPP1, Bradi4g19020.1.v3.1, in blue,
pTM=0.69) and Adiantum nelumboides (AnEPP1, JAKNSL_Ane01050.m, in pink, pTM=0.73), M. paleacea (MpaEPP1, utg000002g0001491 in purple, PTM=0.66). This figure only displays the conserved C-terminal alpha-helixes cluster plus 10 amino acids before the first alpha-helix and 10 amino acids after the
last alpha-helix. The corresponding amino acids are indicated in the figure. Arrows indicate the conserved serine (shown in red) in the loop between the last and penultimate alpha-helixes of the cluster.

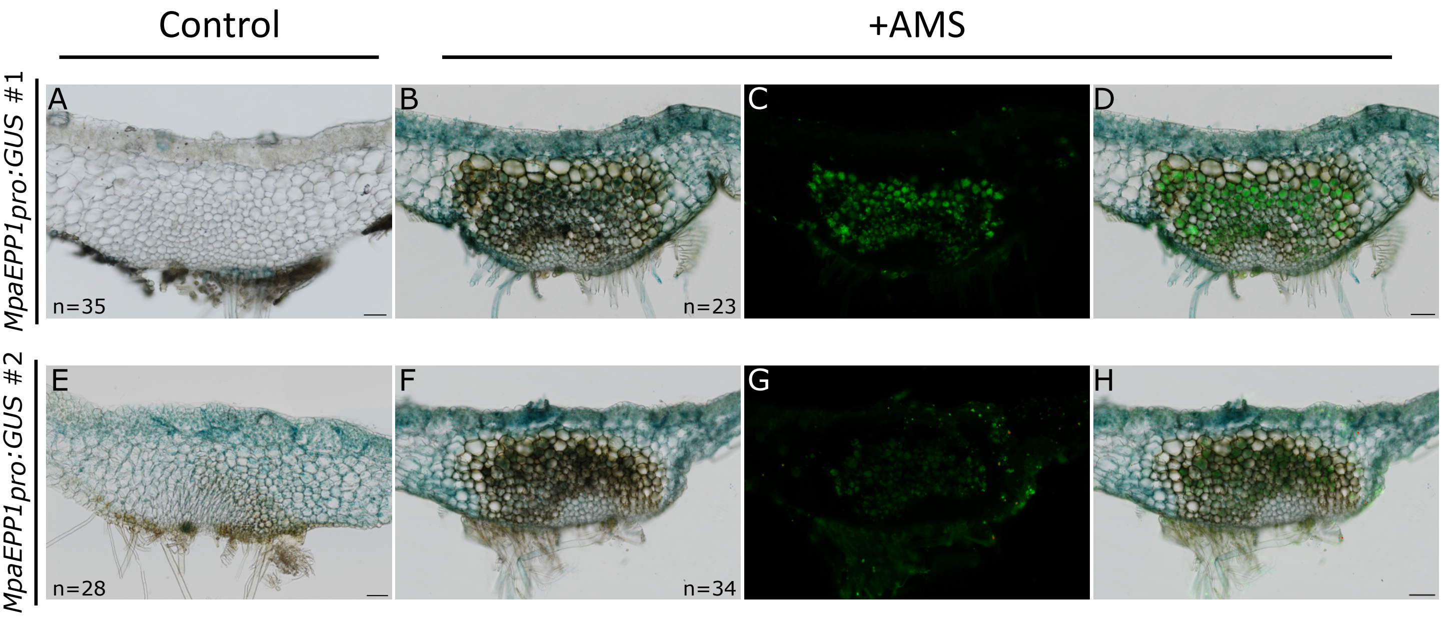


**Supplementary Figure 6.** Two independent *M. paleacea* transformed lines expressing *MpaEPP1pro:GUS* are shown after staining for GUS activity (Scale bar 1mm) after 6 weeks of inoculation with *R. irregularis* (AM,**B,C,D,F,G,H**) or without inoculation (control, **A, E**). Bright field (**A, B, E, F**), Alexa Fluor 488 (**C, G**), and overlays (**D,H**) are shown for each line. Scale bar: 100µm (Nikon microscopy). Total number of thalli observed are indicated.

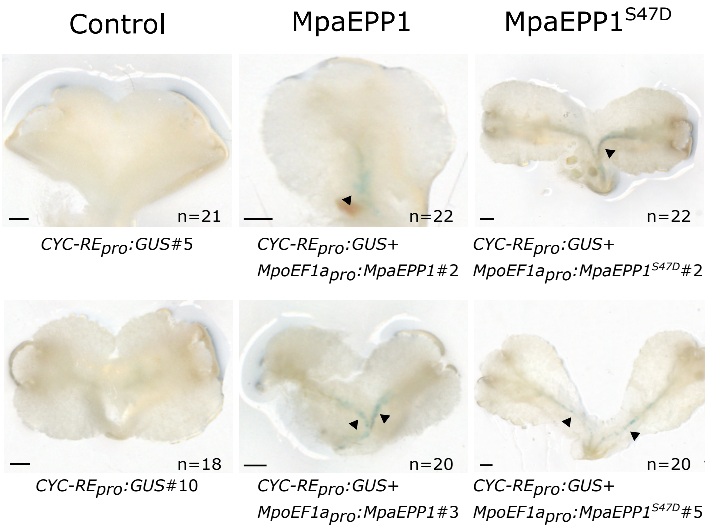


**Supplementary Figure 7.**

*M. paleacea* transformed lines expressing *CYC- RE*$_{pro:}$*GUS* (two independent lines: *CYC- RE*$_{pro}$*:GUS* #5 and 10) or *CYC- RE*$_{pro}$*:GUS* + *MpoEF1apro:MpaEPP1* (two independent lines: #2, 3) or *CYC- RE*$_{pro}$*:GUS* + *MpoEF1apro:MpaEPP1*$^{S47D}$ (two independent lines: #2, 5) are shown after staining for GUS activity. (Scale bar, 1 mm.) Total number of thalli observed are indicated.

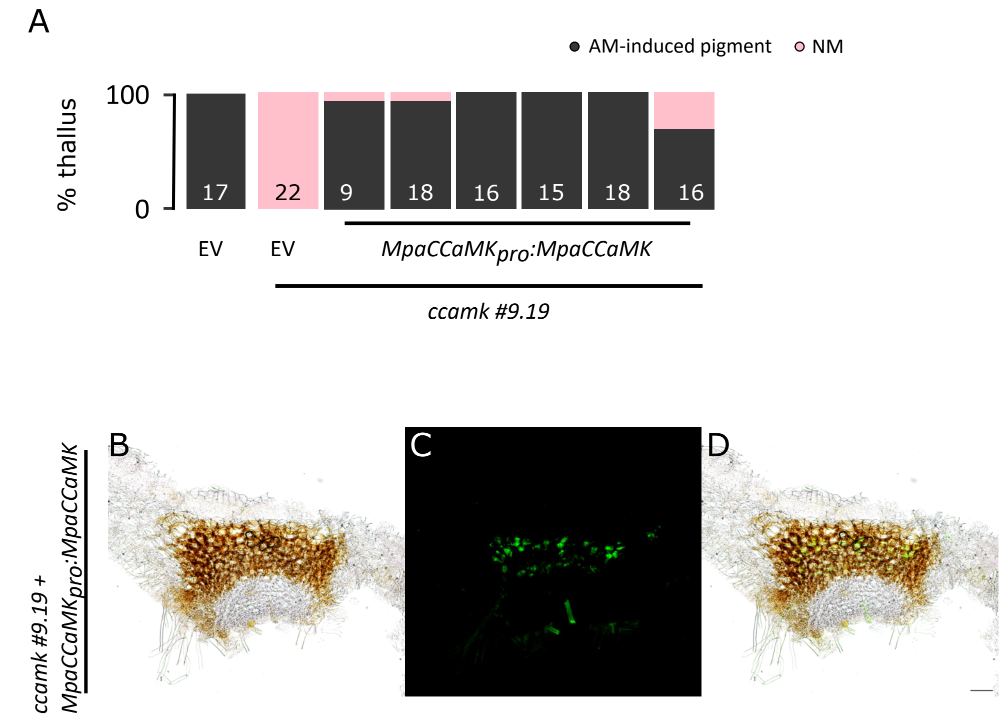


**Supplementary Figure 8.**

**A** Percentage of colonized thalli in wild type plants transformed with en empty vector (EV) or *ccamk #9.19* (Vernié *et al.* 2025) plants complemented with MpaCCaMK under the native *MpaCCaMK* promoter or an empry vector (EV). Number of observed thalli are indicated. Each bar corresponds to an independent transformed line. **B-D** transversal sections of *ccamk #9.19* line transformed with MpaCCAMK under *MpaCCaMK* native promoter six weeks post inoculation with *R. irregularis*. *R. irregularis* is visualized with WGA -Alexa Fluor 488. Bright field (**B**), Alexa Fluor 488 (**C**), and overlay (**D**) are shown for one representative line. Scale bar: 100µm (Nikon microscopy).
